# Development of a high-throughput bead based assay system to measure HIV-1 specific immune signatures in clinical samples

**DOI:** 10.1101/195362

**Authors:** Thomas Liechti, Claus Kadelka, Hanna Ebner, Nikolas Friedrich, Roger D. Kouyos, Huldrych F. Günthard, Alexandra Trkola

## Abstract

The monitoring and assessment of a broadly neutralizing antibody (bnAb) based HIV-1 vaccine require detailed measurements of HIV-1 binding antibody responses to support the detection of correlates of protection. Here we describe the development of a flexible, high-throughput microsphere based multiplex assay system that allows monitoring complex binding antibody signatures. Studying a panel of 13 HIV-1 antigens in a parallel assessment of different IgG subclasses (IgG1, IgG2 and IgG3) we demonstrate the potential of our strategy. The technical advances we describe include means to improve antigen reactivity using directed neutravidin-biotin immobilization of antigens and biotin saturation to reduce background. A particular emphasis of our study was to provide tools for the assessment of reproducibility and stability of the assay system and strategies to control for variations allowing the application in high-throughput assays, where reliability of single measurements needs to be guaranteed.

## 1. Introduction

Broadly neutralizing antibody (bnAb) responses are considered a key element of protective vaccines against HIV-1^1^. Evaluation of vaccine candidates in animal models and in human trials not only requires measuring of the desired function, a broad neutralization activity, but also a thorough assessment of a broad range of binding antibody properties elicited by the vaccine to confirm correlates of protection. This was highlighted by the RV144 study^2,3^, where non-neutralizing antibodies directed against the V2 region of the envelope protein were linked with the modest protection observed in this trial. Besides the observation that these V2 antibodies can induce ADCC, strong IgG3 responses were observed in RV144 highlighting a potential protective effect of Ab effector functions (e.g. antibody dependent cellular cytotoxicity) in this trial^2^. Several other studies reported on the high variability of HIV-1 binding antibody response in terms of IgG subclass distribution, elicitation of effector functions and epitope recognition in natural infection and upon vaccination^2,4,5^.

These types of studies depend on sensitive and accurate binding assays that allow assessment of many parameters in 100s – 1000s of samples. Here we report on the development of a microsphere bead-based multiplex immunoassay technology to measure binding responses against multiple HIV-1 derived proteins and peptides of different Ig subtypes. The method we describe allows improved coupling to beads over published protocols and provides a full quality assessment that allows to run large sample sets without the necessity to obtain repeat measurements. The latter is a pre-requisite for true high-throughput screening assays.

## 2. Methods

### 2.1 Clinical samples and ethics information

All analyzed plasma samples were derived from specimen stored in the biobanks of the Swiss HIV Cohort study (SHCS) and the Zurich Primary HIV Infection Study (ZPHI). The Swiss HIV Cohort Study (SHCS) is a prospective, nationwide, longitudinal, non-interventional, observational, clinic-based cohort with semi-annual visits and blood collections, enrolling all HIV-infected adults living in Switzerland^6^. The SHCS is registered under the Swiss National Science longitudinal platform: http://www.snf.ch/en/funding/ programmes/longitudinal-studies/Pages/default.aspx#Currently%20supported%20longitudinal%20studies. Detailed information on the study is openly available on http://www.shcs.ch. The Zurich Primary HIV Infection study (ZPHI) is an ongoing, observational, non-randomized, single center cohort founded in 2002 that specifically enrolls patients with documented acute or recent primary HIV-1 infection (www.clinicaltrials.gov; ID NCT00537966)^7^.

The SHCS and the ZPHI have been approved by the ethics committee of the participating institutions (Kantonale Ethikkommission Bern, Ethikkommission des Kantons St. Gallen, Comite departemental d'ethique des specialites medicales et de medicine communataire et de premier recours, Kantonale Ethikkommission ZÜrich, Repubblica e Cantone Ticino - Comitato Ethico Cantonale, Commission cantonale d'étique de la recherche sur l'être humain, Ethikkommission beider Basel for the SHCS and Kantonale Ethikkommission ZÜrich for the ZPHI) and written informed consent had been obtained from all participants.

### 2.2 HIV-1 antigens and antibodies

We selected 13 HIV-1 antigens covering structural Gag proteins (p17 and p24) and various Env antigens (gp120 JR-FL, BG505 gp140, BG505 trimer, V3 and MPER peptides, CD4bs probes (RSC3, RSC3Δ) for the development of the assays. Properties, sources and details to production and purification of HIV-1 antigens are listed in **Supplementary Table 1**. We thank contributors to the NIH AIDS Reagent Program, D. Burton (TSRI, USA), J.P. Moore (Cornell University, USA), R. Sanders (AMC, Netherlands), P. D. Kwong (NIH-VRC, USA), M. Nussenzweig (The Rockefeller University, USA), J. Robinson (UZH, Switzerland) for providing proteins, peptides or expression plasmids for this study. All peptides were chemically biotinylated as indicated in **Supplementary Table 1**. With the exception of p24 and gp41ΔMPER all other proteins were expressed with an avi-tag and enzymatically mono-biotinylated (Avidity, LLC, Colorado, USA). **Supplementary Table 2** contains information on all the monoclonal antibodies used in this study.

### 2.3 Detection of HIV-specific IgG isotypes using a customized multivariate multiplex bead assay

To measure plasma IgG1, IgG2, IgG3 binding antibody reactivity to the 13 selected Gag and Env HIV-1 antigens we established a bead-based multiplexed immunoassay using the Luminex technology^©^.Carboxylated MagPlex^©^ beads (Luminex) were either covalently directly coupled with antigens or coupled with Neutravidin (Sigma Aldrich) followed by loading with biotinylated antigens. MagPlex^©^ beads contain unique ratios of two fluorescent dyes which allow the distinction of up to 500 different so-called bead regions. Choosing unique bead regions for each antigen/analyte allowed to perform individual measurements in one simultaneous reaction. Coupling was done using a coupling kit (BioRad) according to the manufacturer's instructions. Briefly, 12.5 Mio beads were sonicated and thoroughly washed with the activation buffer provided by the coupling kit. The carboxyl groups on the beads were activated in 100µl activation buffer containing sulfo-N-hydroxysulfosuccinimide (S-NHS, Thermo Fisher Scientific) and 1-ethyl-3-(3-dimethylaminopropyl)carbodiimide hydrochloride (EDC, Thermo Fisher Scientific) at a concentration of 5mg/ml. After 20 minutes of activation at room temperature beads were thoroughly washed with activation buffer and beads were coupled in 500µl PBS containing the respective proteins (50µg of gp41ΔMPER, 62.5µg p24) for direct coupling or 62.5 µg Neutravidin for 2 hours at room temperature. Beads were then washed thoroughly, blocked with PBS-BSA 1% for 30 minutes at room temperature and after extensive washing stored in PBS-BSA 1% at a bead concentration of 20'000 beads/µl at 4°C. Biotinylated proteins were coated on neutravidin-coupled beads and subsequently blocked with biotin. Proteins and biotin were diluted in PBS at a concentration of 320nM (if not stated otherwise) and coating and blocking were performed for 1hour at room temperature. Beads were extensively washed with PBS-BSA 1% and stored at a concentration of 10'000 beads/µl at 4°C and were used for up to 56 days.

Patient plasma was heat inactivated at 56°C for 1hour before use the binding assay. The different bead regions carrying the different antigens were mixed at each experiment and incubated with patient plasma or monoclonal Abs diluted in PBS-BSA 1% over night at 4°C to allow plasma IgG binding to antigen-coated beads. Beads were washed with PBS-BSA 1% and split up equally into 3 different reactions to allow for detection of IgG1, IgG2 and IgG3 from the same plasma sample. IgG responses were detected with Phycoerythrin (PE)-labeled secondary antibodies specific to isotypes IgG1, IgG2, IgG3 or total anti-human IgG diluted in PBS-BSA 1% at a concentration of 1µ/ml (Southern Biotech) for 1hour at room temperature. After extensive washing with PBS/BSA 1% beads were analyzed with the FlexMap 3D or LX200 instruments (Luminex). A minimum of 50 beads per antigen and plasma were acquired to guarantee accurate MFI values. Quality control and validation procedures for the FlexMap 3D instrument were done on each day of experiment according to manufacturer's instructions.

### 2.4 Detection and exclusion of measurements affected by the Hook effect

To assess the performance of the assay we probed plasma samples from 8 chronically HIV-1 infected individuals against the complete antigen panel. Plasma samples were probed over a range of dilutions, in four completely independent replicates on multiple day. For each plasma dilution curve (where each data point is the geometric mean of the four replicate measurements), we determined the dilution at which the maximal signal intensity was measured (**Supplementary Figure 6**). This is usually the lowest (1:100) dilution. If this was not the case, all lower dilutions were identified as affected by the Hook effect, which is a well-described phenomenon likely caused by an overabundance of plasma antibodies^8^. In **Figure 4** and **Supplementary Figure 7**, where the focus is on the day post bead coupling, we used a stringent approach and excluded, for each antigen and each plasma, all measurements at a fixed dilution where at least the measurement on one of the five days was affected by the Hook effect. This particular choice of exclusion ascertains that all investigated distributions of log binding titers are based on the same plasma samples and measured at the same dilution, while minimizing the number of excluded plasma samples.

### 2.5 Exponential curve fitting

For each patient plasma, each dilution and each antigen, we used an exponential decay model to quantify the observed decrease in signal intensity over time post bead coupling (**Figure 4B**). Let y be the measured MFI and let x be the day post bead coupling. Fitting a linear regression model log_10_ *y* = *m x + b* to the data and transformation of the slope m yields

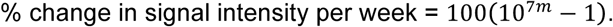

### Assay variability

To assess the performance of the assay we probed plasma samples from 8 chronically HIV-1 infected individuals against the complete antigen panel. Plasma samples were probed over a range of dilutions, in four completely independent replicates on multiple day. We measured the (intra-plasma) assay variability as the coefficient of variation of the log binding titers across the four replicates (**Figure 5**) and as the standard deviation of the log binding titers across the four replicates (**Figure 6**). In addition, we measured the (inter-plasma) assay variability as the standard deviation of the average log binding titers (over the four replicates) across the eight tested plasma samples (**Figure 6**).

### 2.7 Strategy to optimize the choice of dilution(s)

In high-throughput experiments, plasma samples can only be tested at a few dilutions. The following strategy identifies the optimal dilutions for high-throughput experiments given that m≥1 dilutions can be tested. In a pre-experiment, several randomly chosen plasma samples are tested against all antigens of interest and at various dilutions (if possible many, covering a large range); using e.g. a 96-well plate, one could measure 8 plasma samples at 12 dilutions. Let A be the set of antigens, and let D be the set of tested dilutions. For each antigen *a ∈ A* and each dilution *d ∈ D*, the inter-plasma differences (standard deviations of the measurements) are computed, denoted by 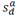.To avoid choosing dilutions that show high inter-patient differences due to the Hook effect, the number of affected measurements is calculated for each dilution and each antigen (see Methods), and 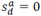 is set manually for antigens and dilutions, where more than 10% (or some even lower threshold) of the plasma samples are affected. The best set *B*^*^ ⊆ *D* of m dilutions is the one that maximizes the sum of the inter-plasma differences,

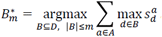

## 3. Results and Discussion

### 3.1 Assay development – oriented, neutravidin-based bead immobilization to improve signal to noise ratio

In the initial phase of our assay development we coupled HIV-1 derived proteins and peptides covalently to the carboxylated beads as described previously^5^ **(Supplementary Figure 1A** and **Supplementary Table 1)**. We used the resurfaced core protein 3 (RSC3)^9^, a protein designed to be preferentially recognized by CD4 binding site (CD4bs)-specific antibodies and the construct RSC3Δ^9^, which lacks a crucial site required for CD4bs Ab binding, to assess the potentials of the assays. Direct coupling of RSC3 and RSC3Δ yielded the expected results both displaying good binding of the glycan specific Ab 2G12 as described^9^ and no cross-reactivity with the V3-specific 1-79, MPER-specific 2F5 or polyclonal human IgG from healthy donors (Figure 1A). In addition RSC3 showed specific binding of the CD4bs specific monoclonal antibodies b12 and VRC01 whereas RSC3Δ did not. While specificity was therefore as desired, the obtained signals were in general low and did not exceed a median fluorescence intensity (MFI) value of 100 with the Luminex LX200 which has an upper MFI limit of 35'000.

Based on this we next took measures to improve the signal-to-noise ratio. Covalent linkage of proteins to carboxylated proteins via primary and secondary amines can result in random orientation of the protein on the beads resulting in masking of the epitope of interest in a fraction of the proteins^10,11^. As this could be a problem for probes like RSC3, we developed a procedure for covalent linking of neutravidin to beads and coating the neutravidin-coupled beads with site-directed biotinylated proteins **(Supplementary Figure 1B)**. This guarantees that all proteins are displayed in the same (known) orientation on the beads.

NA coupling resulted in dramatically increased signals of CD4bs bnAb b12 and VRC01 binding to RSC3 **(Figure 1A)**. At the same time NA coupling had no effect on binding of the same bnAbs to RSC3Δ nor was 2G12 binding to both proteins altered. Identical binding of 2G12 irrespective of the antigen coupling highlights that NA-coupling did not result in higher quantities of proteins on beads compared to direct-coupling and hence NA-coupling must overcome a negative effect on epitope accessibility that exists in the case of direct coupling **(Figure 1A)**.

We next tested the differentially coupled antigens for their capacity to detect CD4bs specific IgG (i.e. RSC3 reactive) in plasma of six individuals with acute (n=3) and chronic (n=3) HIV-1 infection. **(Figure 1B)**. In line with the known delay in Ab development in HIV-1 infection, we detected no Specific CD4bs Ab activity in the three acutely infected persons. In contrast all three chronically infected cases showed reactivity with RSC3, which was higher for NA-coupled versus the direct coupled beads **(Figure 1B)**. These results strongly emphasize that coating of biotinylated proteins on neutravidin carrying beads results in highly improved signal-to-noise ratio compared to the classical approach.

### 3.2 Assay development – reducing background noise

In a separate line of experiments we sought to find means to reduce background noise. Proteins and peptides contain charged moieties that may enhance aggregation. Theoretically aggregated proteins could pose a problem to the measurement, as they may unspecific ally bind to beads and also potentially could be transferred between beads loaded with different probes. During the actual measurement where beads loaded with different proteins/peptides are present this could contribute to a noise signal. We thus introduced a blocking step with free biotin that was applied immediately after coupling the biotinylated probes to NA on the beads. This blocking step ensures occupation of free binding pockets of neutravidin and resulted in dramatically reduced background noise signals as shown for the reactivity to NA-coupled signals with the background control composed of neutravidin-coupled beads without coated biotinylated protein **(Supplementary Figure 2)**.

**Figure 1:**
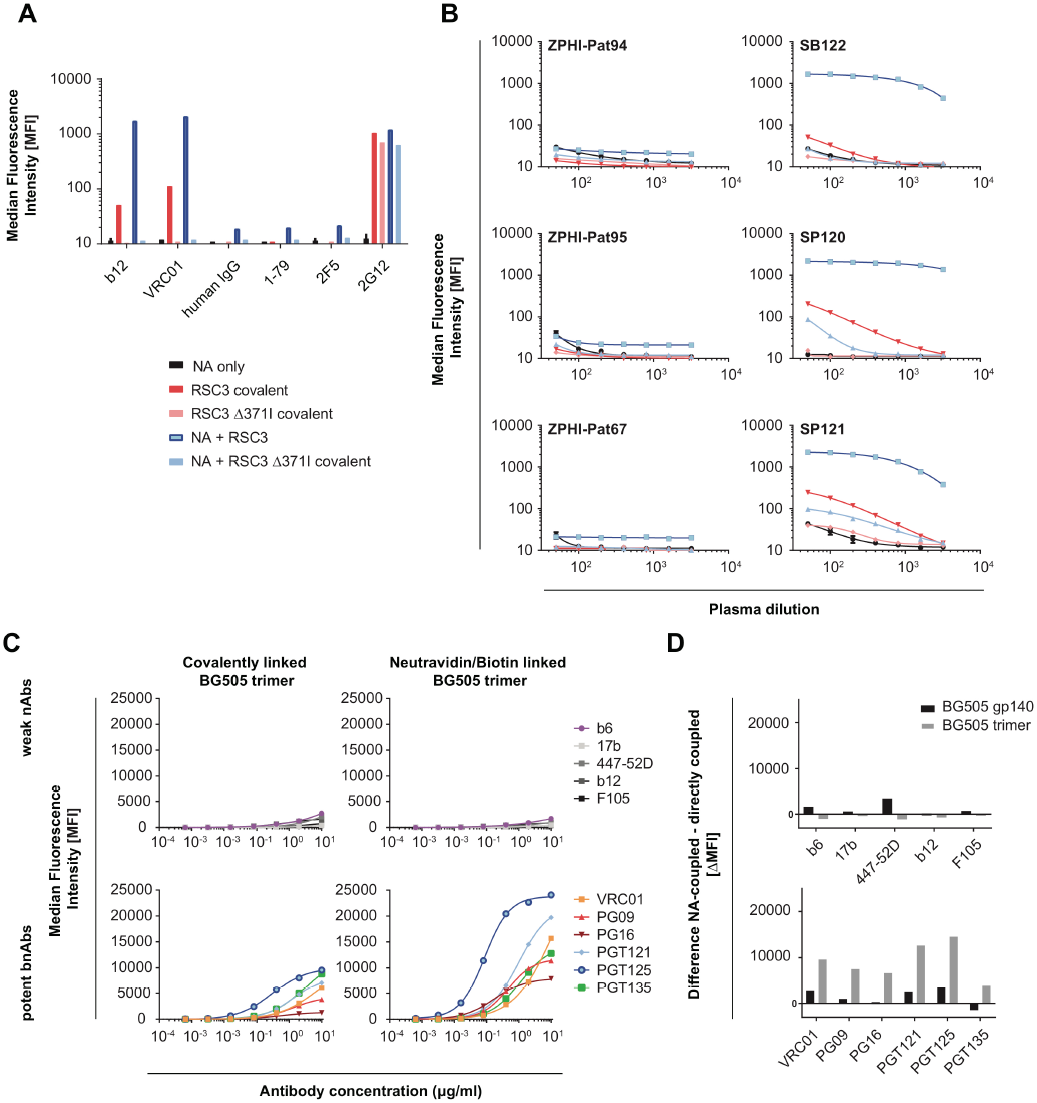
Improving signal-to-noise ratio by developing a new luminex assay based on biotinylated proteins and peptides and neutravidin-coupled beads. (A) Comparison of antibody binding to RSC3 (CD4bs probe) and RSC3Δ (CD4bs deficient probe) coupled covalently to Luminex beads (red bars) or via biotin to neutravidin (NA)-coupled Luminex beads (blue bars) in a Luminex bead binding assay. CD4bs specific antibodies (VRC01 and b12) known to bind RSC3 but not RSC3Δ and the glycan specific nAb 2G12 known to binds both constructs were used to compare signal-to-noise ratio in the two different coupling setups. NA only signifies beads that were not coupled with antigen to establish the background. Purified human IgG from healthy donors, V3-specific mAb 1-79 and the MPER-specific bnAb 2F5 were used as negative controls. (B) Comparison of plasma antibody reactivity of acute (n=3) and chronically (n=3) HIV-1 infected individuals with the direct and NA coupled beads described in A. (C) Comparison of reactivity of gp120-specific mAbs with BG505 trimer coupled covalently to beads (left) or through biotin on NA-coupled beads (right) were tested. Reactivity of weakly neutralizing (top row) and broadly neutralizing Abs (bottom row) are shown. (A-C) Median Fluorescence intensity (MFI) values obtained in a representative experiment on an LX200 reader are shown. (D) The difference in binding signal to BG505 trimer and BG505 gp140 between NA-coupled and directly coupled beads shown in C) is depicted.

### 3.3 Assay development – assessing the advantage of NA-coupled antigens for HIV-1 Ab analysis

To verify if biotinylated proteins provide a general advantage in signal detection, we next assessed binding of several mAbs to BG505 trimer. For this comparison BG505 trimer was either directly coupled to beads or N-terminal biotinylated BG505 was coupled to NA coated beads. The BG505 trimer adopts a closed conformation and thus is known to bind bnAbs more efficiently than non-or weak neutralizing antibodies^12^. Indicators for a closed conformation are bnAbs like PG09 and PG16, which preferentially bind quaternary structures compared to Env monomers. Our comparison of the two coupling methods revealed that irrespective of the coupling method weak-or non-neutralizing antibodies did not bind BG505 trimer (**Figure 1C).** In contrast however, binding of bnAbs to the NA-coupled trimer was greatly increased. **(Figure 1C and D).** Interestingly the increase in PG09 and PG16 was proportionally higher (**Figure 1D**) emphasizing that the biotin/neutravidin approach provides a better preservation of the trimer structure.

The BG505 trimer contains the immunodominant HR1 and HR2 regions of gp41 monoclonal antibodies against the gp41 failed to bind the trimer due to its closed native-like structure^12^. To verify that NA coupling does not lead to a distortion of the conformation in this area, we tested the reactivity of gp41 mAbs with NA-immobilized N-terminally biotinylated trimer. To this end we tested a panel of gp120-and gp41-specific antibodies for the ability to bind to the BG505 trimer or the trimeric gp41 lacking MPER (gp41ΔMPER). The BG505 trimer was only recognized by gp120-directed but not gp41 antibodies. In line with published reactivities^13^, the bnAbs VRC01, PG09, PG16, PGT121 and 2G12 were the most efficient in binding the trimer **(Figure 2A and B)**. While the gp41-specific antibodies did not bind the BG505 trimer, they yielded high signals using the gp41ΔMPER **(Figure 2A and B)**. To further verify this, we tested binding of plasma from 25 chronically infected patients to BG505 trimer and gp41ΔMPER and probed if binding is affected by competition with soluble gp41ΔMPER **(Supplementary Figure 3)**. We observed a dose-dependent decrease in plasma Ab binding to gp41ΔMPER but not the BG505 trimer confirming that gp41-directed antibodies cannot bind to the native-like BG505 trimer. Collectively these results highlight that neutravidin/biotin coupling of BG505 trimer to beads does not alter the conformation of the protein and preserves its intact native-like closed structure.

In a further step, to ensure that our assay provides the best possible signal-to-noise ratio we titrated different concentrations of biotinylated, monomeric gp120 JR-FL on neutravidin-coupled beads to test whether the signal intensity can be improved when the protein density on beads is increased **(Supplementary Figure 4)**. Increasing concentrations of gp120 JR-FL during coating did not result in relevant increases of signals and MFI at saturation and EC50 titers remained largely identical **(Supplementary Figure 4B)** suggesting that at the concentration range used all accessible NA sites are occupied. We therefore set the concentration of proteins during NA-bead coating at 320nM. Similar results were obtained with gp41ΔMPER coupled directly to the beads suggesting that also for direct coupling a wide range of protein concentrations on the beads results in consistent fluorescence intensities **(Supplementary Figure 4C and D)**.

### 3.4 Assay development – probing large NA-coupled bead panels

In a next development step we probed a multiplex panel of 13 HIV-1 protein/peptide targets **(Figure 3).** Eleven of the targets were immobilized via the NA-strategy; the remaining two (p24 and gp41ΔMPER, which were not available with N-terminal biotinylation) were directly, covalently coupled to the beads. Since both p24 and gp41 are highly immunodominant and generally elicit high antibody titers in HIV-1 infection^14,15^, potentially decreased signals due to covalent linkage were not a concern. Using this panel of 13 HIV-1 probes we generated binding titers against all antigens for a range of HIV-1 mAbs with known epitope specificity and a polyclonoal HIV-1 IgG pool from infected patients (HIVIG) **(Figure 3)**. The results show the expected binding pattern for mAbs according to their known epitope specificity. The polyclonal HIVIG bound as expected for HIV-1 plasma antibodies well to p17, p24 and gp41ΔMPER, but had surprisingly weak reactivity with gp120 JR-FL and beyond this with no other gp120 probe suggesting that the overall gp120 Ab content in this pool is low. Collectively, these data verify that the Luminex based multiplex assay we developed is highly specific and that no crosstalk between the targets occurs.

**Figure 2:**
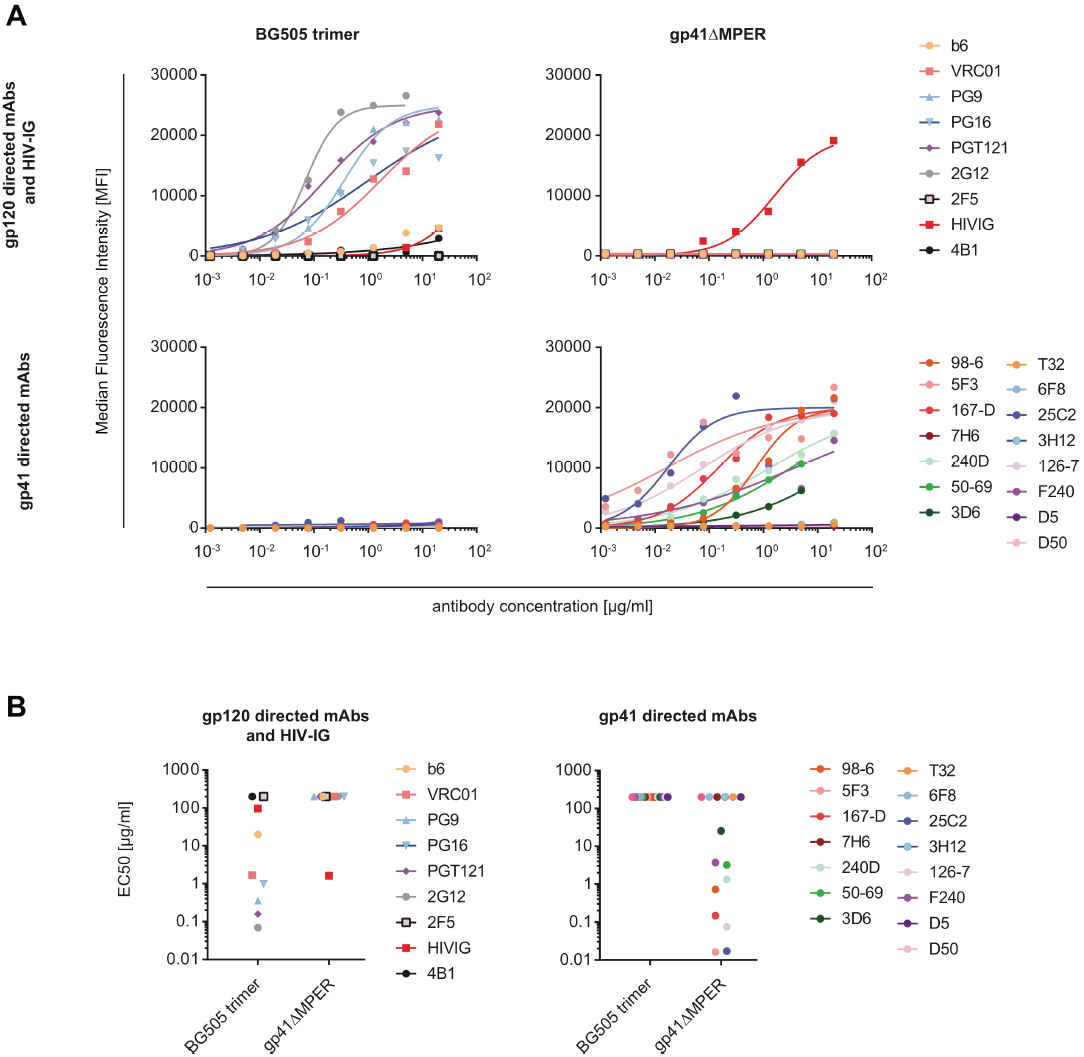
BG505 trimer maintains closed conformation after coupling via biotin on neutravidin-coupled beads. (A) Binding reactivities of gp120-specific (top row) and gp41-specific (bottom row) antibodies against BG505 trimer (left column) and gp41ΔMPER (right column) were compared for directly and biotin/NA-coupled beads in the Luminex binding assay. Median fluorescence intensity (MFI) values obtained in a representative experiment on an LX200 reader are shown. (B) 50% effective concentration (EC50) values derived from data shown in (A) are depicted.

### 3.5 Assay development – increasing the dimensionality of the measurements by parallel assessment of IgG subclasses

A specific goal in our assay development was to create an assay that provides the possibility to not only increase the panel size if this becomes of interest but also to add a further layers of information by assessing specific properties of antibodies. A primary objective for us was to estimate the contribution of IgG1, IgG2 and IgG3 to the HIV-1 binding antibody response. To allow this we defined a strategy where a given plasma sample is first incubated with the full panel of beads. After the IgG binding step, beads are split into three different reactions with specific secondary antibodies against the different IgG subclasses. In order to guarantee that the recorded signals measured are indeed specific for the corresponding IgG isotype, we probed the specificity of the secondary antibodies **(Supplementary Figure 1C and 5)**. For this, commercially available standards for IgG1, IgG2 and IgG3 were incubated with anti-IgG coupled Luminex beads and tested whether the secondary antibodies cross-react with the different IgG subclasses. Our results indicate that the secondary antibodies are specific to the individual subclasses. We saw, however, some degree of cross reactivity of anti IgG-1 detection Ab with the IgG3 standard, although only at very high concentrations **(Supplementary Figure 5)**. Concentration of the IgG-1 detection Ab used routinely in our assay were set to 1µg/ml so that no cross-reactivity could inflict our measurements.

**Figure 3:**
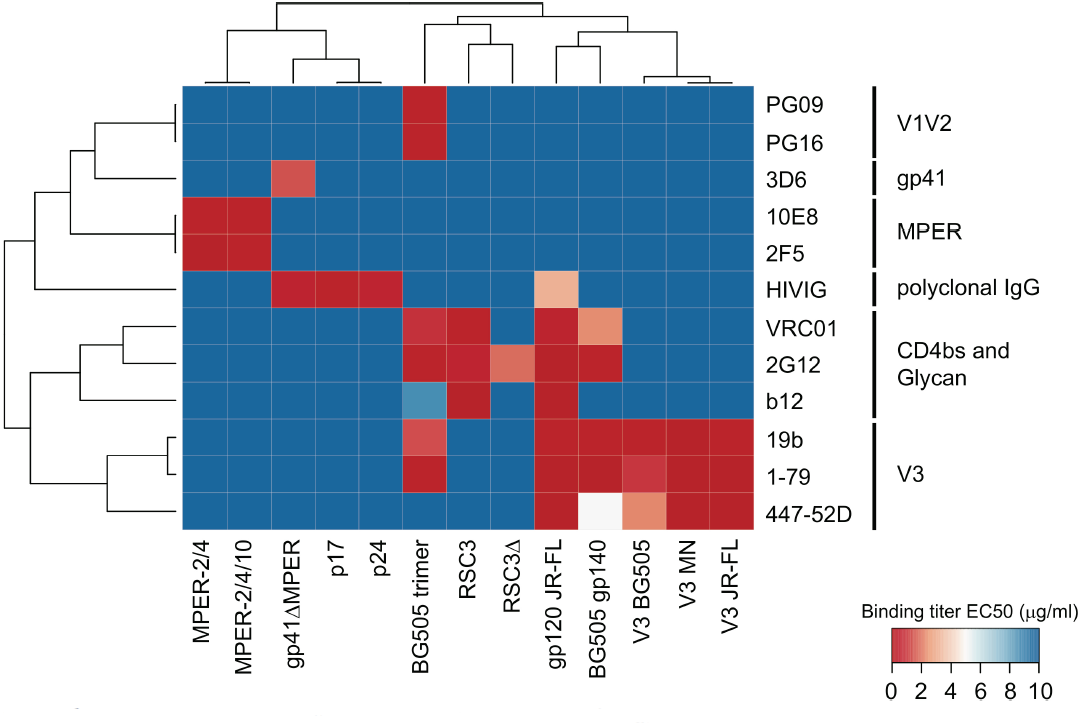
Specificity and accuracy of the multiplex bead binding assay. HIV-1 specific mAbs antibodies with specificities covering all included 13 HIV-1 antigens were tested for reactivity in the multiplex assay. A heatmap of the binding reactivity (50% effective concentration (EC50) values) shows the specificity to mAbs and potential collinearities between proteins/peptides. Hierarchical clustering with the complete linkage method and the Euclidean distance measure was used to cluster antibodies and targets.

### 3.6 Assessment of assay performance

Luminex binding assays have been used in several studies to measure antibody binding reactivities to HIV-1 antigens^5,16,17^ but the ability to detect a broad range of epitopes and the different isotypes contributing to the signal was not yet assessed in detail.

To test the reproducibility, variability of our assay system we conducted an in-depth multiplex binding analysis of plasma samples from 8 chronically HIV-1 infected individuals using the full panel of 13 HIV-1 antigen described above.. The binding reactivities to the 13 HIV-1 antigens were measured on 5 days (1, 6, 14, 34, 56 days post bead coupling) at 6 different plasma dilutions (3-fold dilutions from 1:100 to 1:24300), and in 4 replicate assays. This large, multi-dimensional data set allowed us to analyze the temporal assay stability, the assay variability, and the optimal dilution for each antigen.

For each tested HIV-1 antigen, each plasma sample and each day of testing, we obtained dilution curves where each data point is the geometric mean of four replicate measurements **(Supplementary Figure 6**). For some highly reactive antigens, we observed a clear Hook effect^8^: the maximum signal intensity was not reached at the lowest 1:100 dilution but at a higher dilution **(Supplementary Figure 6)**, which is likely caused by an overabundance of plasma antibodies. 23.9% of all measurements were affected by this effect (see Methods) and we excluded them from all further analyses where they might affect the results.

#### 3.6.1 Temporal assay stability

A specific interest of our analysis was to define temporal affects. As high-throughput screenings for which we designed the assay, commonly will require data acquisition over prolonged periods it is important to monitor and control for a potential decay of the antigen-bead batches. Indeed we observed that the signal intensity decreased continuously over time and that this was already noticeable shortly after coupling of the beads **(Figure 4A).** This decrease was observed for most antigens but more pronounced for antigens with weaker reactivity **(Supplementary Figures 6 and 7)**. Although several explanations for this observed loss of signal intensity may apply, likely reasons are constant detachment of the protein coupled to the beads and protein decay. In line with this, the decrease generally followed an exponential decay pattern **(Supplementary Figure 8).** This decay affected all antigens irrespective of the mode of bead immobilizations but was less pronounced for directly coupled antigens in our panel.

Fitting exponential regression models to curves as in **Supplementary Figure 8** revealed widely varying decay rates across the 13 antigens **(Figure 4B)**, with binding titers to antigens like V3 MN and MPER-2/4/10 decreasing by on average more than 20% per week. This clearly demonstrates that measurements obtained on different days post bead coupling cannot be compared without further adjustments. Based on our experience we strongly advocate that monitoring of temporal effects need to be generally incorporated when assessing reactivity over several days to weeks by bead based assays.

The signal intensity decreased very similarly for all 8 patients **(Supplementary Figure 8)**. This is an important and positive finding as it implies that the day post bead coupling – while it strongly affects the absolute signal strength – does not influence the relative ranking of the measured plasma samples. The Luminex assay can thus be conducted at any day post bead coupling and plasma samples measured at the same day can be compared directly in absolute terms. For larger screens with measurements on multiple days, this also provides the opportunity that the relative rankings of the plasma samples can be compared. To minimize the risk of falsifying the data by the transformation to relative ranks, this requires two key conditions. First, the order of the plasma samples must be randomized before the start of the experiment. Second, the throughput of samples must be sufficiently high. We recently conducted a Luminex binding assay with 4,281 plasma samples using the here described multiplex panel where we measured 122-354 samples per day and recorded values in ordinal ranks (Kadelka et al., in preparation), highlighting that processing sufficient sample sizes are possible.

**Figure 4:**
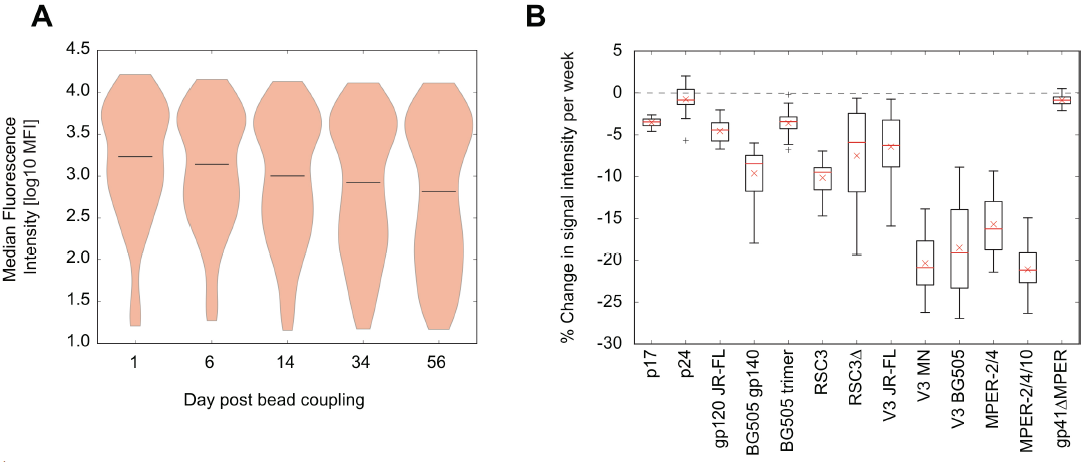
Temporal loss in signal intensity. Evaluation of data depicted in **Supplementary Figure 6**. (A) Distribution of all signal intensities (pooled across all dilutions, all patient plasmas and all antigens) for each day of the experiment, where the signal intensities are the mean over the four replicate assays. For each antigen and each patient plasma, all measurements at a fixed dilution where at least one of the measurements on the five days was affected by the Hook effect were excluded. (B) Distribution of the % change in signal intensity per week post bead coupling for each antigen. For each patient plasma and each non-excluded dilution, the % change values are derived from fitting the data to an exponential curve.

#### 3.6.2 Assay variability

The assay variability, measured as the coefficient of variation of the binding titers across the four replicates, varied strongly (range: 0.0013-0.3425) but was overall small with a median of 0.0457 **(Figure 5A)**. 88.1% of all measurements showed less than 10% variability and even 99.3% less than 20% variability. Immunoassays with a CV of less than 20% are considered accurate^18^,19. Thus the Luminex binding assay we describe here is generally reliable and reproducible.

We conducted further analyses to identify which factors are associated with the assay variability in our assay system. Measurements with a higher signal intensity resulted, on average, in more similar replicates **(Figure 5B).** Accordingly, the variability also differed strongly across antigens with binding to highly reactive antigens being measured more accurately **(Figure 5C).** The choice of plasma dilution also seemed to matter; we observed the lowest average variability for a mid-range plasma dilution of 1:2700 **(Figure 5D).** In addition, the time post bead coupling seemed to influence the variability but the observed pattern was not monotonic **(Figure 5E)**. Importantly, the actual sample measured exhibited no strong effect on the variability highlighting that assay specific factors but not handling or sample composition have an influence **(Figure 5F)**.

### 3.7. Optimizing for high-throughput screens – single measurements at an optimal plasma dilution

In the process of assay evaluations, we measured the binding reactivities at six different plasma dilutions and in four replicate assays. This level of accuracy is not possible in large screens, where to be cost-and time effective analyses ideally need to be restricted to a single measurement of one plasma dilution. It was thus important to probe what the optimal plasma dilution(s) are in our set-up that would allow this. An intrinsic problem of multiplex assays is the highly varying reactivity across different antigens; there is hardly ever a single dilution at which all investigated antigens are measured in the dynamic range. Measurements in the dynamic range of the assay are characterized by comparably higher inter-patient differences, which we exploited to determine the best plasma dilution for each antigen. We used the average standard deviation between the measurements of the 8 patient plasmas as a measure of the inter-patient differences, where the average was taken across the four replicates and the five days of testing. Of the probed dilutions, the dynamic range of highly reactive antigens like p24 or the BG505 trimer seemed best hit by a 1:24'300 dilution, while this dilution was worst suited for measuring RSC3Δ binding as all measurements were in the lower plateau of the assay **(Table 1, Supplementary Figure 6)**. A comparison of the inter-and intra-patient variability of the measurements revealed a significantly larger variation among plasma samples from different donors than among replicates for each antigen (**Figure 6**), which supports the validity of using single measurements in high-throughput experiments.

Above results also show that a range of dilutions yields measurements in the dynamic range of the assay **(Table 1, Supplementary Figure 6)**. Since high-throughput experiments need to be time-and cost-effective, only one, or at most a few, dilutions can be tested. We therefore designed the following strategy to optimize the choice of dilution(s) for a given number of testable dilutions: In a pre-experiment, several randomly chosen plasma samples are tested against all antigens of interest and at various dilutions (if possible many, covering a large range); using e.g. a 96-well plate, one could measure 8 plasma samples at 12 dilutions. Since the dynamic range of an assay is characterized by large differences among the measured plasma samples, our strategy allows us to choose those dilutions for use in a high-throughput screening that yield the highest sum of inter-plasma differences across all tested antigens (see Methods for details). If at all possible, this strategy ensures that there is at least one dilution for each antigen that yields measurements in the dynamic range of the assay.

**Figure 5:**
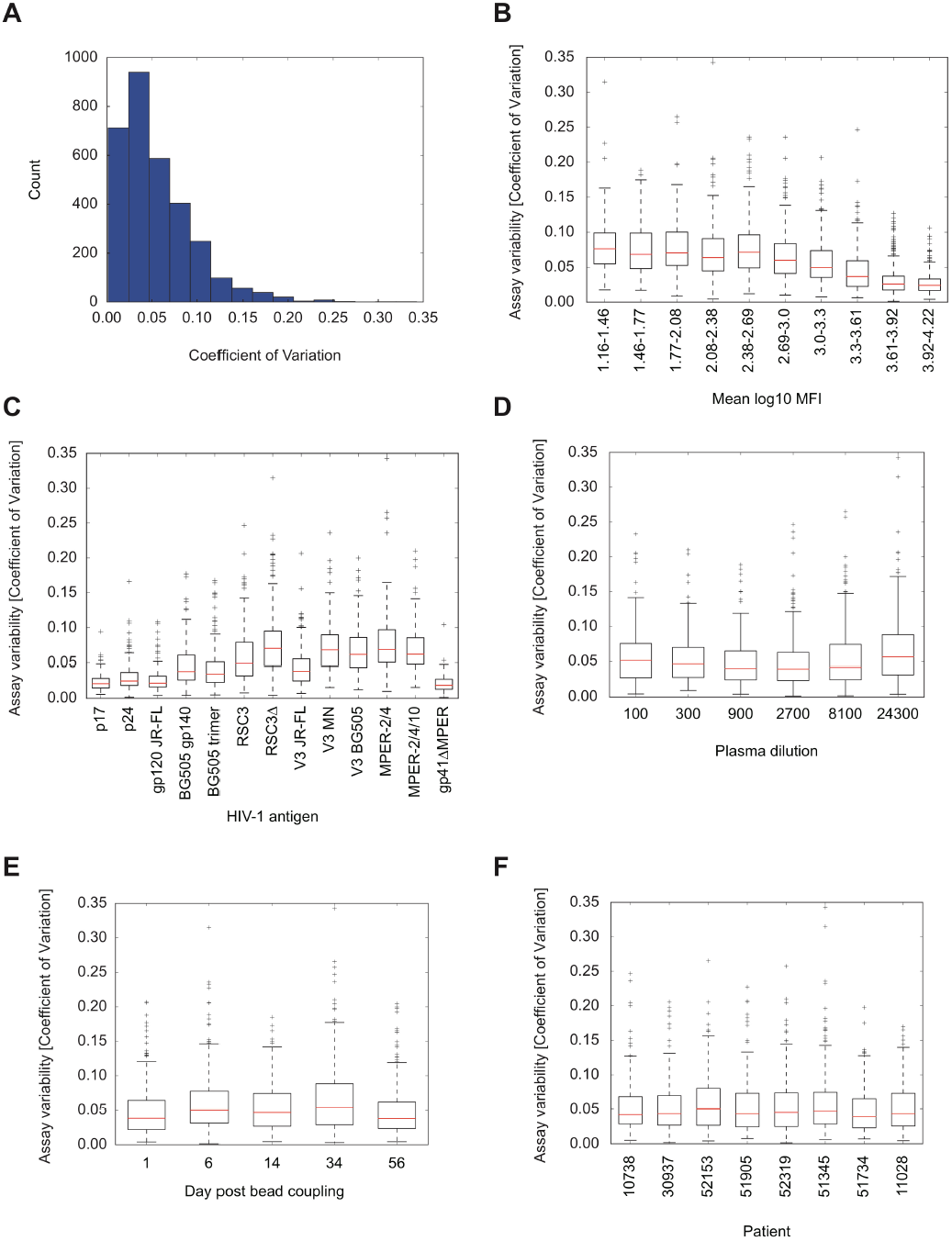
Assessment of assay variability. Evaluation of data depicted in **Supplementary Figure 6**. (A) Histogram of the assay variability. (B-F) Boxplot showing the distribution of the assay variability stratified by mean binding titer (B), antigen (C), dilution (D), day post bead coupling (E), and patient plasma (F). Each red line shows the median assay variability, each box depicts the interquartile range (IQR), and each whisker extends to the most extreme value no more than 1.5*IQR from the box.

**Table 1:** Choice of optimal plasma dilution. Evaluation of data depicted in **Supplementary Figure 6**. (A) The inter-patient differences, measured in standard deviations, are shown for each antigen and each dilution. Standard deviations where more than 10% of the measurements are affected by the Hook effect are written in gray and excluded from the choice of the optimal dilution. Bold font highlights the antigen-dependent optimal dilution, which maximizes the inter-patient differences. (B) Example of the strategy designed to pick best dilutions for high-throughput screens. The sums considered for determining the next best dilution are computed from the values in A, and at each step, the best dilution to be added is highlighted in shades of green.

|  |  | Dilution |  |  |  |  |  |
| --- | --- | --- | --- | --- | --- | --- | --- |
|  |  | 1:100 | 1:300 | 1:900 | 1:2700 | 1:8100 | 1:24300 |
|  |  | Average inter-patient standard deviation |  |  |  |  |  |
| HIV-1 antigens | p17 | 0.098 | 0.052 | 0.026 | 0.021 | 0.033 | **0.068** |
|  | p24 | 0.116 | 0.083 | 0.102 | 0.161 | 0.304 | **0.558** |
|  | gp120 JR-FL | 0.072 | 0.040 | 0.021 | 0.027 | 0.055 | **0.131** |
|  | BG505 gp140 | 0.097 | 0.055 | 0.061 | 0.099 | 0.168 | **0.247** |
|  | BG505 trimer | 0.090 | 0.046 | 0.046 | 0.092 | 0.152 | **0.207** |
|  | RSC3 | 0.101 | 0.056 | 0.070 | 0.162 | 0.357 | **0.529** |
|  | RSC3Δ | 0.344 | 0.417 | **0.421** | 0.353 | 0.214 | 0.063 |
|  | V3 JR-FL | 0.073 | 0.047 | 0.028 | 0.068 | 0.182 | **0.253** |
|  | V3 MN | 0.114 | 0.123 | 0.140 | 0.180 | **0.221** | 0.221 |
|  | V3 BG505 | 0.178 | 0.142 | 0.137 | 0.203 | **0.278** | 0.265 |
|  | MPER-2/4 | 0.218 | 0.259 | 0.316 | 0.368 | **0.375** | 0.302 |
|  | MPER-2/4/10 | 0.198 | 0.212 | 0.222 | 0.230 | **0.231** | 0.184 |
|  | gp41ΔMPER | 0.098 | 0.052 | 0.029 | 0.031 | 0.033 | **0.051** |

**Table 1: Choice of optimal plasma dilution**
|  |  | Dilution |  |  |  |  |  |
| --- | --- | --- | --- | --- | --- | --- | --- |
|  |  | 1:100 | 1:300 | 1:900 | 1:2700 | 1:8100 | 1:24300 |
| **Find best dilution**<br>sum of the average inter-patient standard deviations<br>(excluding Hook-effect affected ones) |  | 0.344 | 0.464 | 1.413 | 1.964 | 2.605 | **3.080** |
| **Find best two dilutions**<br>sum of the gains in average inter-patient standard deviations<br>(excluding Hook-effect affected ones) |  | 0.281 | 0.354 | **0.410** | 0.401 | 0.285 | - |
| **Find best three dilutions**<br>sum of the gains in average inter-patient standard deviations<br>(excluding Hook-effect affected ones) |  | 0 | 0 | - | 0.060 | **0.082** | - |

For our data, the strategy identified the 1:24300 dilution as the best dilution (**Table 1B**). When allowing two dilutions, it added the 1:900 dilution, which provides a large gain in inter-patient differences for RSC3Δ, an antigen that is measured very poorly at the 1:24300 dilution. The two identified dilutions plus the 1:8100 dilution were identified as the best set of three dilutions. The gain in total inter-patient differences per added dilution dropped quickly indicating support for the use of only a few dilutions in high-throughput experiments as the high associated cost of additional dilutions may often outweigh the benefit of a small increase in signal accuracy.

**Figure 6:**
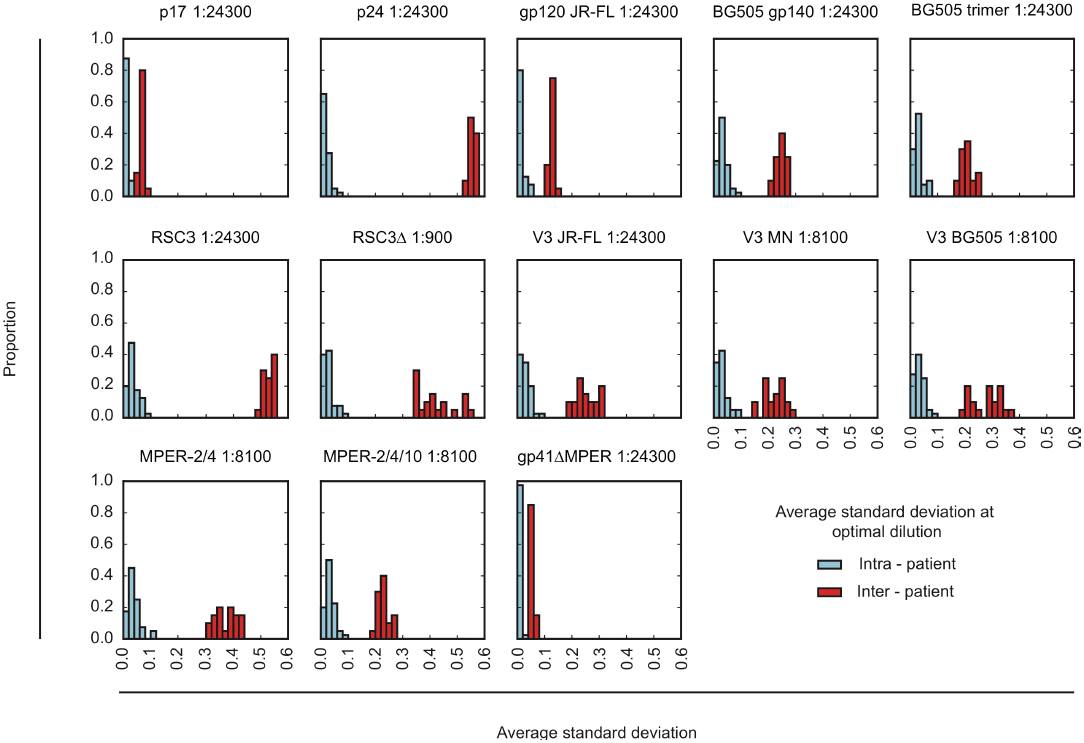
Inter-and intra-patient variability. Evaluation of data depicted in **Supplementary Figure 6**. The distribution of intra-patient variability (average standard deviation of the measurements among the four replicates; blue) and inter-patient variability (average standard deviation of the measurements among the eight patients; red) is compared for each antigen at its optimal dilution (see **Table 1A**).

## 4. Conclusion

With antibody-based HIV-1 vaccine development moving closer to large-scale HIV vaccine trials the necessity to have accurate assay systems that allow high-throughput and high-content analysis increases. Here we describe the development and validation of a sensitive multiplex bead-based assay system that allows for simultaneous detection of antibodies binding distinct target sites on the envelope trimer and other HIV-1 derived antigens. The assay system provides in its current setup means to differentiate HIV-1 binding antibody signatures in HIV-1 infected individuals or vaccinees. Using the same procedures as we applied here (NA coupling of targets, validation) other targets or host responses can be easily approached. As we lay out here, a comprehensive knowledge of the assay performance in terms of variability and stability is required to allow application in high-throughput testing. Importantly, we show that once the relevant factors are known, they can be controlled for making accurate high-throughput measurements possible.

## Acknowledgements

Financial support for this study has been provided by the Swiss National Science Foundation (SNF; #314730_152663 and #314730_172790 to AT), the Clinical Priority Research Program of the University of Zurich (Viral infectious diseases: Zurich Primary HIV Infection Study to HFG and AT), the Swiss Vaccine Research Institute (to AT, HFG, RDK) and the SystemsX.ch grant AntibodyX (to AT). RDK was supported by the SNF (#PZ00P3-142411 and BSSGI0_155851). This study has been co-financed within the framework of the Swiss HIV Cohort Study, supported by the SNF (# 33CS30_148522 to HFG), by the small nested SHCS project 744 (to AT) and by the SHCS research foundation. The funders had no role in study design, data collection and analysis, decision to publish, or preparation of the manuscript. The SHCS data are collected by the five Swiss University Hospitals, two Cantonal Hospitals, 15 affiliated hospitals and 36 private physicians (listed in http://www.shcs.ch/180-health-care-providers). We thank the patients participating in the ZPHI and the SHCS and their physicians and study nurses for patient care and Danièle Perraudin and Mirjam Minichiello for administrative assistance. We are very grateful to Jan van Gils, Tomasz Zborowski and the Luminex cooperation for providing the FlexMap 3D instrument for the HIV IgG measurements.

